# Model-Based Estimates for Operant Selection

**DOI:** 10.1101/2022.07.22.501082

**Authors:** Matthias Borgstede, Patrick Anselme

## Abstract

We present a new methodology to partition different sources of behavior change within a selectionist framework based on the Price equation – the Multilevel Model of Behavioral Selection (MLBS). The MLBS provides a theoretical background to describe behavior change in terms of operant selection. Operant selection is formally captured by the covariance based law of effect (CLOE) and accounts for all changes in individual behavior that involve a covariance between behavior and predictors of evolutionary fitness (e.g., food). In this article we show how the CLOE may be applied to different components of operant behavior (e.g., allocation, speed, and accuracy of responding), thereby providing quantitative estimates for various selection effects affecting behavior change using data from a published learning experiment in pigeons.

## 1 Introduction

Operant learning has repeatedly been characterized as a process that works analogous to natural selection (Broadbent, 1961; D. T. Campbell, 1956; Gilbert, 1970; Herrnstein, 1964; Palmer & Donahoe, 1992; Pringle, 1951; Skinner, 1966; Thorndike, 1900). Whereas in natural selection, species adapt to the environment as a result of the fitness consequences of inheritable traits, operant selection consists of individuals adapting to specific contexts as a result of the consequences of repeatable actions. Skinner (1981) proposed that both processes should be subsumed under the common explanatory mode of *selection by consequences*.

Although the conceptual framework of selection by consequences has a long tradition in behavior analysis and remains a popular narrative (e.g., Baum, 2023; Becker, 2019; Donahoe, 2011; Donahoe, Burgos, & Palmer, 1993; Hull, Langman & Glenn, 2001; Simon & Hessen, 2019; Simon, 2020), it seems to have had little effects on the actual practices of many behavior analysts. One possible reason for this gap between theory and practice may be that operant selection is sometimes used as a mere synonym for what is traditionally called “reinforcement.” Of course, adopting the language and vocabulary of evolutionary biology alone does not add much to the theoretical foundations of the experimental analysis of behavior. To be useful for the development of substantive theory, the conceptual framework of operant selection needs to be scrutinized and formalized, such that it becomes more than a loose analogy. Given such a formal account of operant selection, methodological implications may be derived that might eventually affect the way behavior analysts frame their experiments.

In this article, our goal is to provide a first building block for a methodology of behavior analysis that builds on the conceptual framework of operant selection. In particular, we aim to estimate the amount of selection in different components of operant behavior. To reach this objective, we provide a coherent theoretical background for operant behavior that builds on the concept of selection by consequences (Skinner, 1981). We start with an introduction to the selectionist account of operant behavior and its formalization within the Multilevel Model of Behavioral Selection (MLBS) (Borgstede & Eggert, 2021). Second, we use the MLBS to develop a methodological approach that allows for the empirical estimation of operant selection by means of theory-based modeling. Third, we apply our methodology to derive the amount of operant selection on different components of behavior using training data from a published pigeon experiment. Finally, we discuss the implications of our approach with regard to behavioral selection theory and potential practical applications of the method.

## 2 Behavioral Selection Theory

The principle of operant selection was initially formalized by Baum (2017) and further developed by Borgstede and Eggert (2021), who integrated individual-level behavioral selection with population-level natural selection within the MLBS. The MLBS builds on the abstract description of selection processes by means of the Price equation (Price, 1970, 1972). The Price equation describes selection as the result of the covariance between the individual values of a quantitative character (e.g., an inheritable trait, such as size) and individual evolutionary fitness (i.e., the contribution of an individual to the future population). A positive covariance is associated with a positive change in mean character value (i.e., selection results in a higher average character value), whereas a negative covariance indicates a negative change (i.e., selection results in a lower average character value). In other words, those character values that are statistically related to evolutionary fitness, are selected and thus alter the population average.

In the Price equation, all other sources of change (i.e., those that do not refer to selection) are subsumed in a residual term. This term has different interpretations depending on the context. For example, when applied to the evolutionary change of gene frequencies, the residual term captures the effects of imperfect transmission (i.e., mutation and recombination). When applied to phenotypic change over generations (e.g., body weight or behavioral traits), the term may capture environmental factors influencing the phenotype (Luque & Baravalle, 2021). The Price equation provides a mathematical description of all selection processes, without relying on the specific mechanisms of variation, selection, and transmission (Luque, 2017). In fact, in its most general form, the partitioning of change into a selection and a non-selection term is merely a mathematical identity. Therefore, it is more of a formal definition of what is meant by *selection*, rather than a statement about hypothetical mechanisms. The benefit of such a definition is that it provides a consistent conceptual background that can then be used to construct more specific models of evolutionary change.

In the MLBS, the Price equation framework is applied to a population of individuals that vary in a certain quantitative behavior (such as average number of pecks emitted in the presence of certain environmental cues). At the population level, the covariance term refers to the effects of natural selection on average behavior tendencies of the population, and the residual term captures the average change within individuals. The main contribution of the MLBS is that it formally links this latter within-individual change to the general framework of the Price equation. Following the rationale that individual changes in behavior can also be explained through selection by consequences, the MLBS extends the Price equation by applying the same covariance principle at the within-individual level. Here, the population average in behavior is not calculated over different individuals, but over recurring instances of the same context (e.g., experimental trials). Behavior change, such as an increase or decrease in key pecking rate, averaged over a longer period, is described according to the covariance partitioning from the Price equation. However, the criterion of selection is not individual fitness itself (in terms of a direct contribution to the future population), but statistical predictors of individual fitness. In other words, at the within-individual level, behavior is not selected by means of reproduction or survival (in fact, if the individual dies, all of its behavior immediately ceases), but by events that predict expected change in evolutionary fitness (Borgstede, 2020; 2024). For example, food is generally a positive predictor of evolutionary fitness because it raises the probability of survival and, thus, future reproduction. Conversely, physical threat is a negative predictor of evolutionary fitness because it lowers the chances of survival and future reproduction. The concept of a fitness predictor is largely equivalent to what Baum (2012) calls a “phylogenetically important event” (PIE).

The conceptual framework of the MLBS allows us to describe behavioral selection at the individual level by means of a within-individual covariance between behavior in recurring contexts and its consequences in terms of statistical fitness predictors.^1^ Similar to the process of natural selection, those behaviors that covary with events that signal a change in expected evolutionary fitness are selected, which in turn changes the average behavior of the individual. Following the MLBS, the amount of behavior change due to selection corresponds to the covariance between behavior and fitness predictor, weighted by the slope of the fitness function of the fitness predictor. Formally, the amount of behavior change due to within-individual selection can be expressed by the following equation (Borgstede & Eggert, 2021):

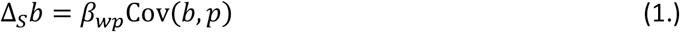

Here, *b* designates a quantitative behavior (such as the amount of key pecking in a recurring context) and Δ_*S*_*b* corresponds to the within-individual change in average behavior due to selection. The value *p* is a quantitative fitness predictor and Cov(*b, p*) is the covariance between the behavior and the fitness predictor. Finally, *β*_*wp*_ is the slope of the function relating the fitness predictor *p* to the actual fitness *w*.

Just like the original Price equation, the MLBS introduces a non-selection term that captures all other sources of behavior change. Designating the overall change in average behavior as Δ*b*, the within-individual behavior change can now be expressed as the sum of a selection term and a non-selection term *δ*:

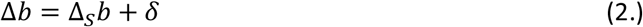

This equation is called the covariance based law of effect (CLOE).^2^ The CLOE can be regarded as a fundamental principle of behavior analysis in that it captures the essence of behavioral selection by partitioning the overall change in behavior into a selection component and a non-selection component (Borgstede & Luque, 2021). The selection component largely corresponds to what is traditionally called reinforcement (i.e., contingency-based effects of behavioral consequences), whereas the non-selection term subsumes all other sources of behavior. Note that Equation 2 is valid for any combination of fitness predictors and behavioral measures. Therefore, instead of absolute counts of pecking and food delivery, relative amounts of pecking of behavior over two options (“allocation”) may be used. Likewise, additional behavioral parameters may equally be treated as potential targets of selection, such as peck frequency (“speed”) or the average success rate of pecking (“accuracy”). In the following section, we use the MLBS as a starting point for a theory-based approach to quantifying selection as a source of within-individual behavior change.

## 3 Fundamental Principles, Models, and Measurement

At a theoretical level, the MLBS accounts for a great range of behavioral phenomena that are difficult to explain using traditional stimulus-response theories (cf. Borgstede, 2021; Borgstede & Eggert, 2021; Borgstede & Luque, 2021), such as the blocking effect (Kamin, 1969), operant selection by means of rare behavior (Premack & Premack, 1963), the establishment of effective contingencies through response deprivation (Timberlake & Allison, 1974), the relation between operant selection and information seeking (Berlyne, 1957; Hendry, 1965), and various deviations from matching (Davison & McCarthy, 2016).

However, it is less obvious how the MLBS might be applied to actual empirical data from behavioral experiments. The reason for this gap between theoretical explanation and empirical application lies in the nature of the Price equation. Since the Price equation is a mathematical identity, it makes no empirically testable predictions per se. Consequently, a fundamental theoretical principle such as the CLOE cannot itself be put to empirical test. The CLOE, like all fundamental theoretical principles, is best understood as a formalization of the conceptual framework used in the underlying theory. In other words, the CLOE tells us what exactly is meant by operant selection and, in doing so, provides the conceptual groundwork for more specific models that may then be applied to empirical data (cf. Borgstede & Luque, 2021; Killeen, 2023).

Although the idea that a fundamental theoretical principle itself has no empirical content may seem at odds with the foundations of empirical science, it is in fact the rule rather than the exception. For example, some of the most fundamental “laws” of behavior are actually true by definition (cf. Killeen, 1962). The same holds for fundamental principles in other sciences, such as physics. For example, Newton’s second law of motion (*F* = *ma*) alone says nothing about any particular physical system. It is only through the construction of specific models by means of auxiliary assumptions and empirically derived regularities that concrete applications become possible (cf. Borgstede & Eggert, 2023). In other words, fundamental laws provide the theoretical backbone of more specific models that may then be applied to actual empirical data. Some of these applications may serve as critical experiments in the evaluation of the theory as a whole. Others may exploit the theory by estimating hitherto unknown model parameters. Such latter applications often do not question the theory itself, but use it to infer the specific values of one or more theoretical entities in a given context. A well-known example in behavior analysis consists in using the generalized matching law to estimate the bias and sensitivity parameters in a given context (cf. Baum, 1974). Applications that seek to infer the values of theoretical entities may not only provide useful practical information, but also form the foundation of theory-based measurement (Borgstede & Eggert, 2023). The aim of the following section is to provide a corresponding methodology for the estimation of operant selection from behavioral experiments.

## 4 Model-Based Inference of Operant Selection

Although the partitioning of change into selection and non-selection components by means of the Price equation is always possible at a theoretical level, empirical applications require specific models for the dynamics of change. Estimation of selection effects becomes possible by comparing observed data to the predictions from such models. The basic rationale behind this approach consists of constraining a specific model such that it predicts what would be expected in the absence of selection and contrasting this prediction with the actual empirical observations. Consequently, an empirical application of the MLBS requires a model of the specific conditions that generated the observed behavioral data. In particular, the model needs to account for the covariance between a behavior and a fitness predictor in a given context (e.g., a specific behavioral experiment).

As outlined in Borgstede and Luque (2021), the theoretical covariance between an observed behavior and a quantitative fitness predictor can be obtained from the feedback function of a reinforcement schedule. Moreover, the quantitative behavior itself (and, consequently, behavior change) can be observed over several repeated experimental trials. Given the feedback function of the target behavior in a specified context, the CLOE may then be applied to empirical data. If the aim is to obtain quantitative values for selection, one can use the model to calculate the amount of operant selection as defined in the MLBS. Technically, selection estimates are calculated using a constrained model (or *null-model*) that is identical to the model that describes the observed behavior, except for the part that is responsible for selection to occur (cf. Okasha & Otsuka, 2020). Practically, this means to apply a minimal change to the model parameters, such that the covariance term becomes zero (because a zero covariance implies zero selection).

Mathematically, there are several ways to ensure that the covariance term in the CLOE is zero. However, the most plausible candidate for our null-model is certainly that the fitness predictor (e.g., the amount of food received per time) is set equal across trials. For example, if we conduct an experiment with two consecutive trials, we may observe behavior that yields three food items per minute in trial 1 and five food items per minute in trial 2. The null-model would use the feedback function from trial 2 to calculate the behavior that would have resulted given the individual had received the exact same amount of food items per minute as in trial 1 (i.e., only three food items per minute instead of five). Conditional on the MLBS, the difference between the actually observed behavior in trial 2 and the behavior predicted from the null-model corresponds to the amount of behavior change that can be ascribed to selection. The corresponding selection estimate may be calculated for any component of the observed behavior, such as relative time allocation, peck frequency, or average success rate of pecking, the only difference being that the feedback function used in the null-model needs to be specified such that it captures the effects of the corresponding target behavior. The proposed method can thus be summarized by the following steps. First, describing the experimental scenario in terms of behavioral selection using a specific model that is consistent with the MLBS. Second, constructing a null-model to calculate the amount of behavior change that would have occurred in the absence of selection. Third, subtracting the behavior change predicted from the null-model from the actually observed behavior change.

In the following section, we will demonstrate the method outlined above by applying it to an actual empirical data set. We will show how empirical estimates of selection for various components of operant behavior can be obtained, and how these model-based estimates can be evaluated, compared and tested for statistical significance using a permutation test framework.

## 5 Application: Operant Selection Between Learning Trials

We demonstrate how the general methodology described above may be implemented in an empirical study using a minimal example for illustrative purposes. We apply the method to the data from two training trials (first and last days) of a published behavioral experiment involving pigeons (Anselme et al., 2022). The focus of the main experiment was to investigate the effects of differential distribution of food items per patch in the holes of a board on foraging behavior. Here, we focus on the training trials administered prior to the main experiment, consisting of two conditions only (“no food” vs. “guaranteed food” at the beginning of a trial). The question we address is to which extent several components of the pigeons’ behavior may have been the target of operant selection and whether the estimated selection effects are significantly distinct from zero.

### 5.1 Experimental Apparatus and Data Acquisition

We exploited some video data from a published study (Anselme et al., 2022) to obtain suitable data for the application of the method outlined above. Here, we only provide the methodological details that are relevant to understanding our analyses.

Nine naïve pigeons were maintained at 85-90% of their free-feeding body weight to motivate them to eat in the task. The pigeons were tested in a rectangular wooden box whose floor was a horizontally removable plate of wood (120 cm length × 70 cm width × 2 cm height), perforated with holes (1.5 cm diameter and ±1.5 cm depth). The foraging board contained 60 holes organized as 6 rows of 10 holes regularly spaced. The board was covered with a black plastic tape with a cross cut above each hole to create an opening, which allowed the pigeons to access the food items while being unable to visually detect their presence from a distance (Figure 1). Specific stimuli (green and red; 21 cm length × 14.5 cm width) were used to signal the consistent presence or absence of food per hole in one area. The two areas were separated by means of a colored strip glued on the plastic tape, dividing the board in two equal left and right areas of 30 holes each from the entrance compartment. Each session was recorded with an external camera, placed above the apparatus.

**Figure 1.**
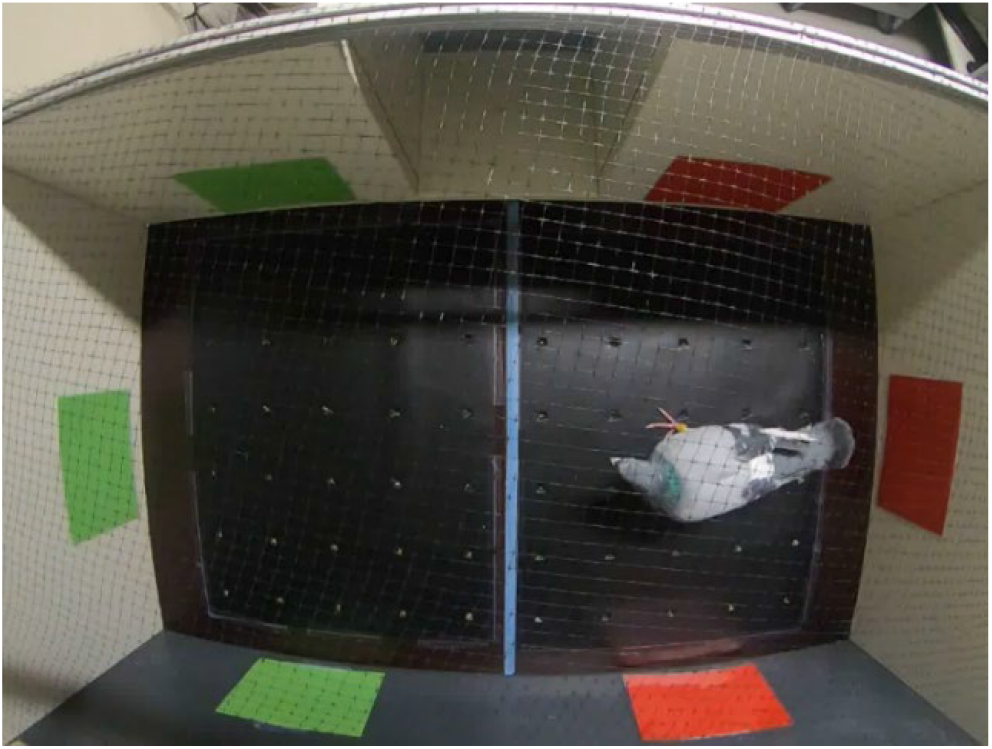
Experimental setting. Pigeons were put in a 120cmx70cm experimental chamber with 60 covered holes. In the food region, each hole contained a grain. In the no-food region, the holes were empty.

In each of the 30 holes of one area (left or right, counterbalanced across trials within the same individuals), we positioned one food item (corn, green pea, yellow pea, or sunflower) and this area was associated with one discriminative stimulus (red or green, counterbalanced across individuals) placed on each wall (4 cm above the floor level), from the first to the last trial. For a given individual, a stimulus location was counterbalanced across trials. The 30 holes of the adjacent area remained empty and were associated with the other stimulus placed on each wall. Of note, the pigeons were initially trained for 3-4 days with uncovered holes, each containing one grain, such that they could see the food in the holes on inspection. No discriminative stimulus was used at this stage. After this initial training, the pigeons were trained as reported above for 4 consecutive days with covered holes.

#### Data Extraction

Data were collected on manual counting (food items consumed and number of pecks per area for each trial). Determining whether a peck at a hole was successful (food item grasped) or not was mostly impossible from the videos, so that peck is not synonymous with item consumed. A peck simply means a vertical downshift of the pigeon’s head above a hole. We considered a pigeon to be positioned in a given area if its head was in this area—because its body could be in one area and yet pecking in the adjacent one. Sometimes, the pigeon missed a grain (picked it up and lost it), so that it rolled on the board. In the attempt to get it, the pigeon could cross the demarcation line between the two areas. A peck given outside of a hole, even to get a missed grain, was not counted.

### 5.2 Data Analysis

We focused on the observed quantitative changes in individual foraging behavior between trials and its relation to individual capture rates. The primary measures used in the analysis were the time spent at the food and the no-food region (*T*^+^ and *T*^−^, respectively), the number of pecks emitted while staying at the food region and the no-food region (*B*^+^and *B*^−^, respectively), as well as the number of food items retrieved during each trial (*R*). Since the most plausible predictor of evolutionary fitness in the current scenario is the retrieval of food per time (capture rate, *C*), we divided the number of retrieved food items during a trial by the total amount of time spent foraging (i.e., the total time the pigeon spent either in the food or the no-food region during each trial), such that *C* = *R/T* with *T* = *T*^+^ + *T*^−^.

As possible targets of operant selection, we calculated three derived behavioral measures. First, relative time at the food region (time allocation, *A*) was calculated by dividing the time spent at the food region by the total foraging time, i.e., *A* = *T*^+^/*T*. Second, differential peck frequency (peck speed or velocity, *V*) was calculated by dividing the number of observed pecks at the food region by the time spent at the food region for each trial, i.e., *V* = *B*^+^*/T*^+^. Third, the average success rate of a peck emitted at the food region (peck accuracy or skill, *S*) was calculated by dividing the number of retrieved food items by the number of pecks at the food region, i.e., *S* = *R/P*^+^.

Since pigeons are known to forage systematically, thereby avoiding sites that they already exploited (Baum, 1987), the expected number of food items per peck only depends on the average pecking success, yielding a feedback function that is approximately linear until all grains are retrieved (for higher numbers of pecks, the slope of the feedback function is zero).^3^ The slopes of the feedback functions correspond to the average gain in capture rate per unit change in the corresponding behavior for each trial. The average gains follow directly from the definitions of the derived measures. The slopes of the corresponding feedback functions are a result of the equality *C* = *AVS*.^4^ For time allocation, rearrangement of the above identity yields a slope of *β*_*CA*_ = *VS*, for peck speed, the corresponding slope is *β*_*CV*_ = *AS*, and for peck accuracy, we obtain *β*_*CS*_ = *VA*, respectively. As an illustrative example, the linear feedback function for time allocation of one individual (P118) is depicted in Figure 2. The dashed black line depicts the feedback function for the first trial that was obtained from the observed time allocation and capture rate during day 1. The observed data from the first trial are indicated by the filled black circle that lies on the dashed black line. The feedback function and data for the last trial (day 4) are indicated by the dashed black line and another filled black circle, respectively.

**Figure 2.**
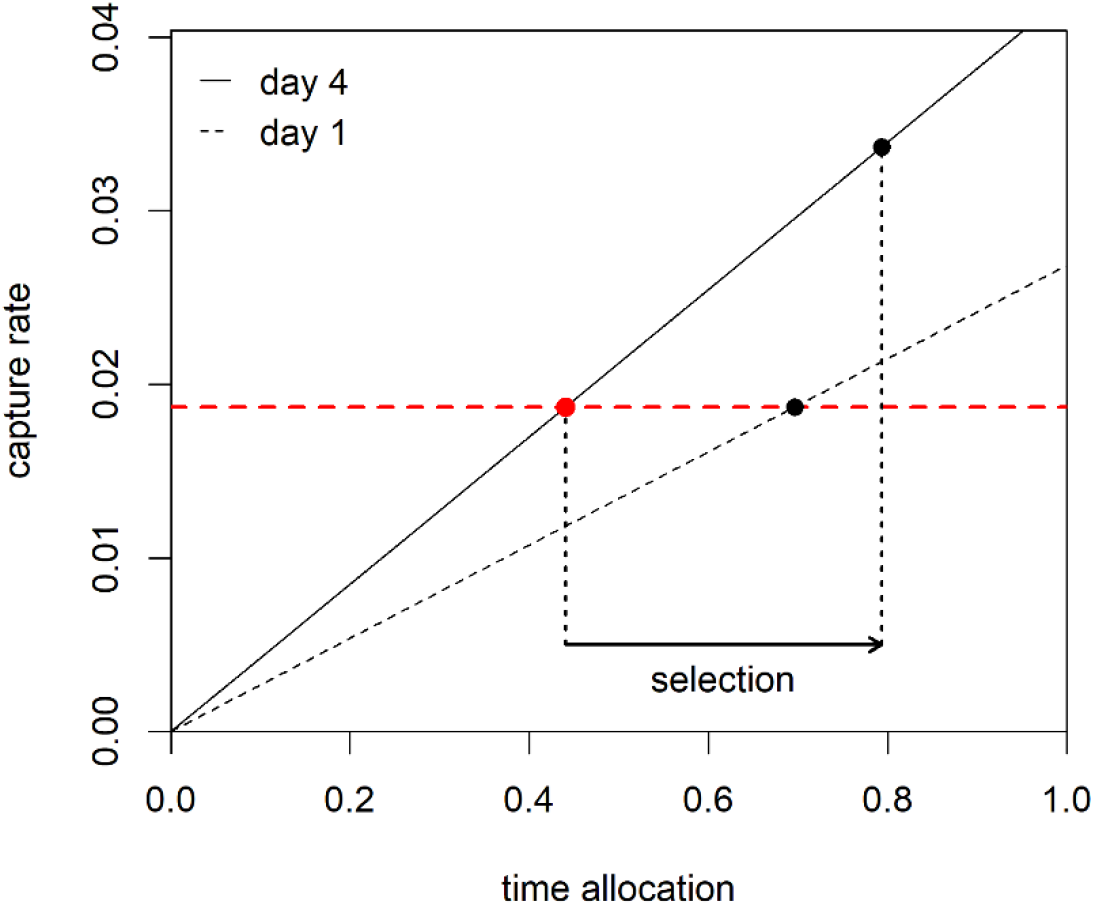
Calculation of selection estimate for time allocation (data obtained from Pigeon 118). The dashed and solid black lines depict the feedback functions for days 1 and 4, respectively. The filled black circles mark the observed behavior (time allocation) and the corresponding feedback (capture rate) for days 1 and 4, respectively. The red dashed horizontal line indicates the observed capture rate during the first trial (day 1). The point where the feedback function of the last trial (day 4) intersects with the red dashed line is marked by a filled red circle and corresponds to the expected behavior given the capture rate had not changed from day 1 to day 4. The difference between this predicted time allocation and the actually observed time allocation in the last trial equals the estimated amount of operant selection acting on time allocation (indicated by the solid black arrow at the bottom of the panel).

The null-model for each behavioral measure was constructed by replacing the observed capture rate on day 4 by the observed capture rate on day 1, thereby constraining the change of the quantitative fitness predictor to zero. In Figure 2, this constraint is illustrated by the horizontal dashed red line that indicates the capture rate on day 1. The point where the red dashed line intersects with the feedback function from day 4 (solid black line) designates the data predicted from the null-model for day 4 (marked with a filled red circle). The corresponding value on the horizontal axis for predicted time allocation on day 4 is then compared to the observed time allocation on day 4. The difference between these two values quantifies the amount of behavior change that can be attributed to operant selection and is indicated by a horizontal arrow in Figure 2. The selection estimates for the other two behavioral measures were calculated analogously using the respective feedback functions for peck speed and peck accuracy.^5^

To test whether the theory-based selection estimates were significantly different from zero, we performed two-sided exact permutation tests (Edgington & Onghena, 2007). We also tested the absolute change values observed for each behavioral measure for significant deviations from zero using two-sided exact permutation tests to evaluate whether the model-based estimates provided any information over and above the raw data. All statistical analyses were conducted using the software R, version 4.2.2 (R Core Team, 2022).

### 5.3 Results

Table 1 summarizes the pigeons’ behavior during the first and the last trial of training, respectively. The average time allocation during the first trial was 0.8 (*SD* = 0.09), indicating that pigeons already spent most of their foraging time at the food region during day 1. The average time allocation during the last trial was 0.76 (*SD* = 0.14). Thus, average time allocation decreased over training trials. The average peck speed at the food region was 0.53 (*SD* = 0.27) on day 1 and was 0.71 (*SD* = 0.28) on day 4, suggesting an increase in mean peck speed. Average peck success also increased from 0.07 (*SD* = 0.05) on day 1 to 0.13 (*SD* = 0.07) on day 4 but was unexpectedly low even after four days of training. Visual inspection of the video material revealed that it often took the pigeons several attempts (sometimes up to 10 pecks or more) to retrieve a food item from a hole in the board. The difficulty of the tasked hence appears to be related to motor skills, rather than failure of food detection. Of the three behavioral measures, the exact paired samples permutation test was only significant for the change in accuracy (*p* = .004).^6^ Average capture rate increased from 0.03 (*SD* = 0.01) on day 1 to 0.06 (*SD* = 0.04) on day 4. The corresponding paired samples permutation test indicated that the observed increase in capture rate was significantly different from zero (*p* = .004).

**Table 1:** Comparison of first and last training trial (day 1 and day 4) with respect to time allocation (time at the food region divided by total foraging time), peck speed (number of pecks divided by time at the food region), peck accuracy (number of retrieved food items divided by number of pecks at the food region), and capture rate (number of retrieved food items divided by total foraging time).

|  | time allocation |  | peck speed |  | peck accuracy |  | capture rate |  |
| --- | --- | --- | --- | --- | --- | --- | --- | --- |
|  | day 1 | day 4 | day 1 | day 4 | day 1 | day 4 | day 1 | day 4 |
| **P17** | 0.95 | 0.96 | 0.40 | 0.15 | 0.09 | 0.26 | 0.03 | 0.04 |
| **P73** | 0.84 | 0.63 | 0.67 | 0.89 | 0.04 | 0.05 | 0.02 | 0.03 |
| **P91** | 0.89 | 0.48 | 0.84 | 0.91 | 0.05 | 0.11 | 0.04 | 0.05 |
| **P96** | 0.74 | 0.80 | 0.81 | 1.11 | 0.05 | 0.15 | 0.03 | 0.14 |
| **P97** | 0.83 | 0.76 | 0.52 | 0.80 | 0.05 | 0.13 | 0.02 | 0.08 |
| **P103** | 0.72 | 0.78 | 0.11 | 0.77 | 0.02 | 0.06 | 0.00 | 0.04 |
| **P118** | 0.70 | 0.79 | 0.39 | 0.63 | 0.07 | 0.07 | 0.02 | 0.03 |
| **P519** | 0.73 | 0.74 | 0.22 | 0.49 | 0.20 | 0.14 | 0.03 | 0.05 |
| **P534** | 0.79 | 0.91 | 0.82 | 0.64 | 0.05 | 0.15 | 0.03 | 0.09 |
| **M** | 0.80 | 0.76 | 0.53 | 0.71 | 0.07 | 0.13 | 0.03 | 0.06 |
| **SD** | 0.09 | 0.14 | 0.27 | 0.28 | 0.05 | 0.07 | 0.01 | 0.04 |

Table 2 shows the amounts of selection on time allocation, peck speed, and peck accuracy that were estimated from the corresponding null-models by calculating the difference between the predicted and the observed values during the last trial (see Section 5.2 and Appendix 2 for details). All three behavioral measures yielded positive selection estimates for all nine pigeons. Average change was largest for time allocation (*M* = 0.38, *SD* = 0.25) and peck speed (*M* = 0.38, *SD* = 0.29), and less expressed for peck accuracy (*M* = 0.06, *SD* = 0.04). However, given that accuracy was very low throughout all sessions, this difference appears to express the overall difficulty of food retrieval, rather than a lower selection pressure. The exact two-sided permutation tests revealed that the selection estimates differed significantly from zero for all three behavioral measures (*p* = .004 for each test). Note, however, that the significance tests are not independent of each other, since the three selection estimates are positively correlated to a considerable degree (correlation coefficients ranging between 0.77 and 0.9). Figure 3 presents a graphical comparison between the mean values and standard deviations for observed change and behavioral selection for time allocation, peck speed, and peck accuracy, which supports the conclusion that selection significantly differs from zero in all three behaviors.

**Table 2:**
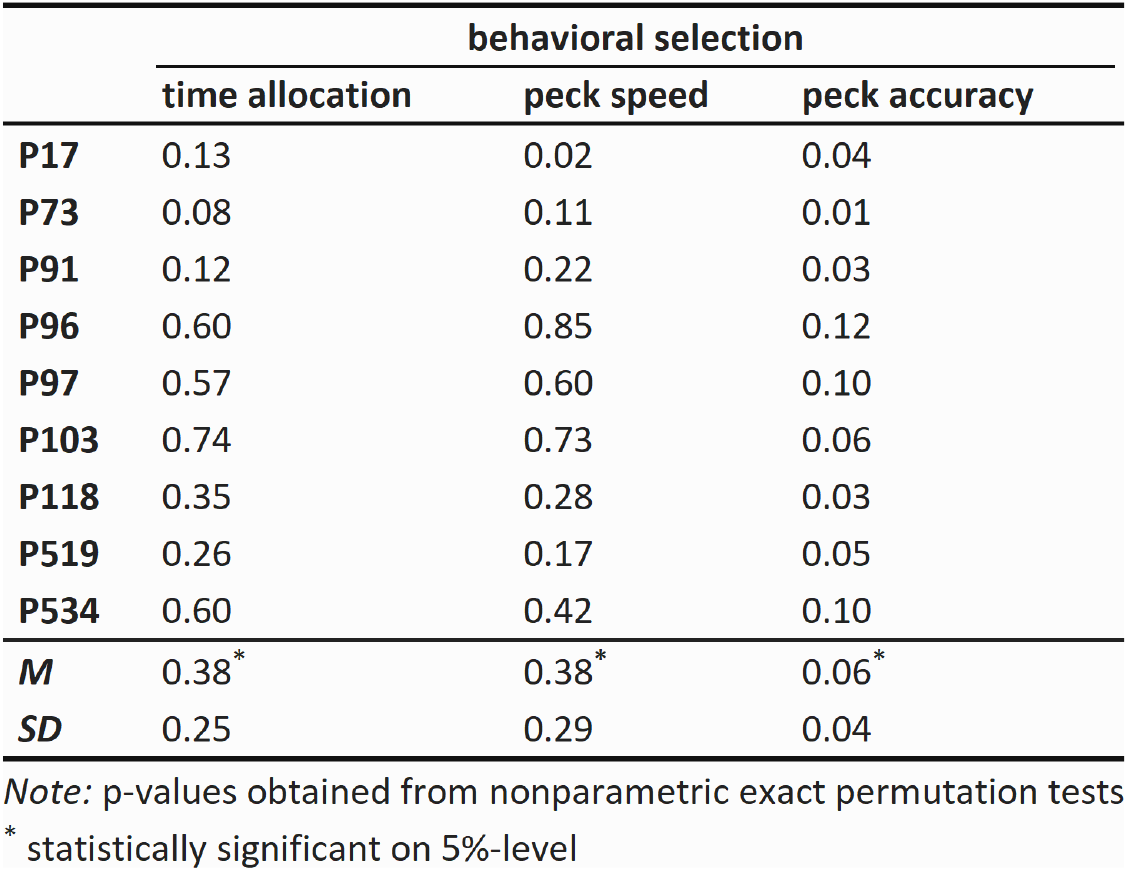
Selection estimates for time allocation (time at the food region divided by total foraging time), peck speed (number of pecks divided by time at the food region), and peck accuracy (number of retrieved food items divided by number of pecks at the food region). The corresponding group-level p-values were obtained from two-sided exact permutation tests.

**Figure 3.**
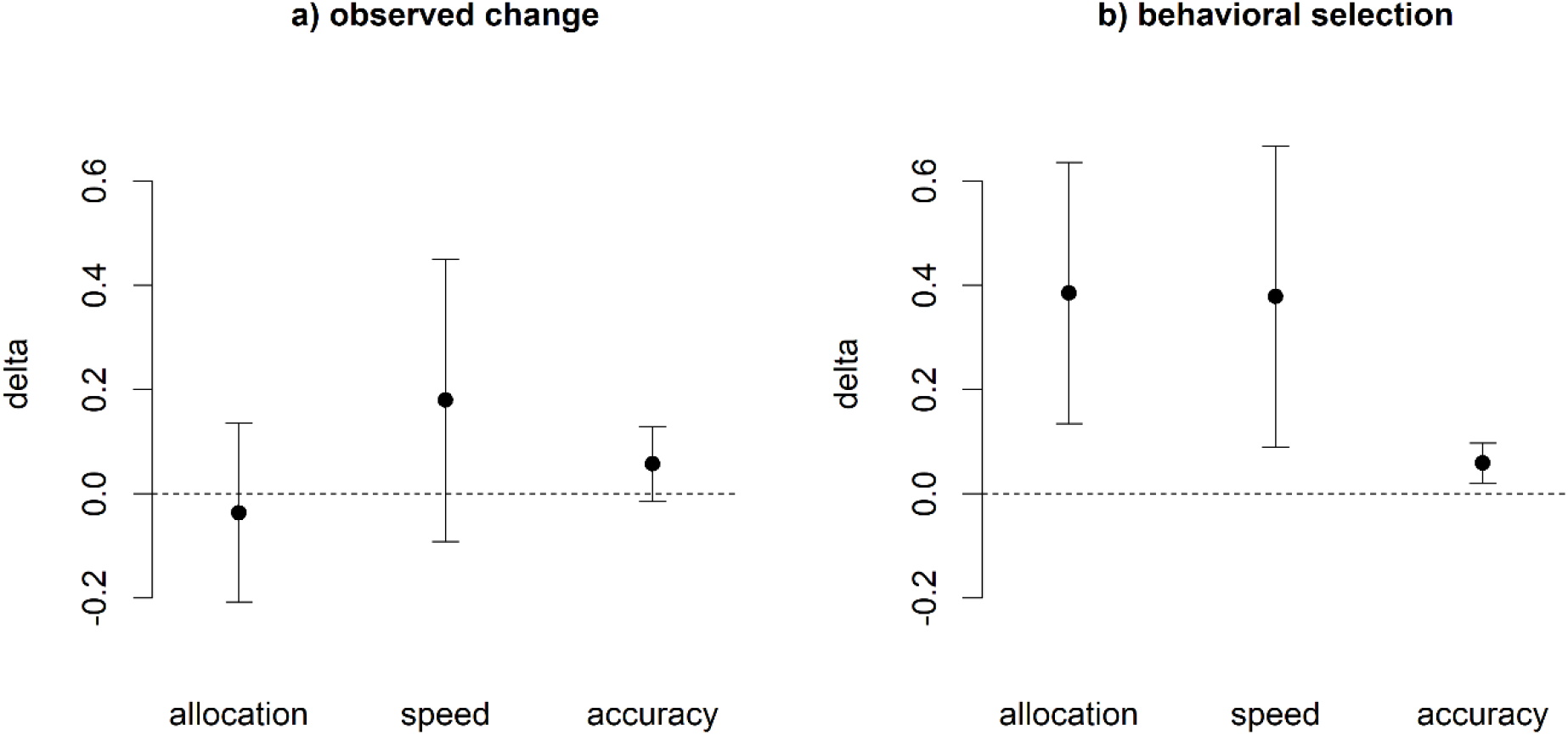
Group means and standard deviations for observed change and behavioral selection, respectively. The vertical axis depicts the change from day 1 to day 4 (delta) for the raw data (panel a) and the model-based selection estimates (panel b). The filled circles indicate the group means, and the error bars the respective standard deviations. Whereas the raw deltas are less expressed with error bars overlapping with zero, the selection estimates are larger on average and clearly differ from zero change.

## 6 Conclusion

In this paper, we proposed a new method to quantify the amount of operant selection in behavioral experiments by means of model-based estimation. The method builds on a formal theory of operant selection, the Multilevel Model of Behavioral Selection (MLBS), which provides an explicit definition of operant selection in terms of an extended Price equation. Applying the MLBS to empirical data, we showed how selection may be inferred for different behavioral measures, such as time allocation, peck speed, and peck accuracy. The rationale was to use the MLBS to construct a null-model that predicts the expected change in behavior in the absence of selection. The difference between the observed behavior change and the predicted behavior change yields an estimate of the selection component of operant behavior.

The method allowed the estimation of different selection effects (i.e., selection on time allocation, peck speed, and peck accuracy) using data from a published foraging experiment. In contrast to the observed raw behavior changes, the selection estimates all significantly differed from zero, indicating that selection was effective even in cases where it was not obvious from the raw data alone. The data further revealed that the selection estimates of allocation, speed, and accuracy were not independent of one another. This latter result is hardly surprising, since the estimation procedure for any of the three behavioral components assumes that the other two behavioral components remain unchanged. However, actual changes in allocation, speed, and accuracy are likely to affect each other. For example, if a pigeon learns where food can be found, this might speed up pecking activity at the relevant area. Therefore, the selection estimates are not to be interpreted as independent additive effects. Instead, they tell us how much change in a certain behavior would be attributable to operant selection if selection was acting exclusively on this behavior.

We presented the first empirical application of the MLBS in the context of a behavioral experiment. The experiment itself was chosen such that it is as simple as possible, to serve as a minimal example for the method proposed in this article. Of course, there are various limitations with regard to the data because they were originally collected for a different purpose. For example, the foraging board was constructed in a way that one could not unequivocally identify the retrieval of food on the videos. Making the floor below the cover transparent might have solved this problem (cf. Baum, 1987). However, this would have possibly enabled the pigeons to see where the food is (because of the light emerging from the holes in the absence of food), a situation likely to affect their foraging behavior. Despite the limitations of the experimental application, our results show that the MLBS in combination with the model-based estimation approach provides a feasible theoretical foundation for the experimental analysis of behavior.

The general methodology of model-based selection analysis can be applied to many other experimental settings that involve behavior change over time. Probably, there are thousands of unused training data sets only awaiting to be analyzed. We hope that this article contributes to the foundations of behavioral selection as a general theory of behavior, and encourages other researchers to put the behavioral selection perspective into practice.

## 7 Author Contributions

MB developed the modeling framework and the statistical methodology, analyzed the data and wrote the original draft. PA planned and conducted the experiments, collected the data and contributed to the theoretical background and the interpretation of the results. Both authors contributed to the final version of the manuscript.

## 9 Data and Code Availability

The original data and R-code used in this study will be made publicly available on publication.

## 11 Appendix

### 11 Derivation of the Covariance Based Law of Effect

The Covariance Based Law of Effect (CLOE) is basically a recursive expansion of the elementary Price equation (Price, 1970):

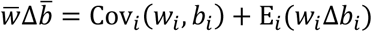

In the Price equation, *w*_*i*_ designates an individual’s evolutionary fitness and *b*_*i*_ the value of an arbitrary evolving character. 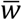 and 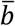 designate the population averages of *w*_*i*_ and *b*_*i*_, respectively. Cov_*i*_ and E_*i*_ are the population covariance and the expected value of the population. Δ*b*_*i*_ is the individual-level change in character value (usually thought of as change between parent and offspring) and 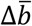 is the population-level change in character value.

The term Cov_*i*_(*w*_*i*_, *b*_*i*_) captures the effects of natural selection, while the term E_*i*_(*w*_*i*_Δ*b*_*i*_) refers to changes in the population average of *b* that are not natural selection. If the time frame is chosen sufficiently small and individuals are treated as their own offspring, E_*i*_(*w*_*i*_ Δ*b*_*i*_) captures changes that occur within individuals.

In the MLBS, the fitness-weighted within-individual change, *w*_*i*_Δ*b*_*i*_, is itself partitioned into a covariance term and an expectation term:

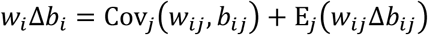

Whereas in the original Price equation, the covariance and expectation are taken over the individuals *i* of a population, in the MLBS, they are taken over a collection of recurring contexts *j* (so-called behavioral episodes) that are themselves nested within individuals. Consequently, the covariance term Cov_*j*_(*w*_*ij*_, *b*_*ij*_) refers to the part of within-individual change that can be attributed to selection at the individual level (i.e., reinforcement), whereas the expectation term E_*j*_(*w*_*ij*_Δ*b*_*ij*_) captures all sources of within-individual change that are not selection.

Given an arbitrary fitness predictor *p* (e.g., food), evolutionary fitness can be predicted by a linear regression of the form *ww* = *β*_*wP*_*p* + *ε*. We can now re-arrange to obtain:

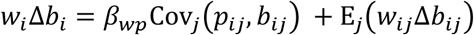

Defining 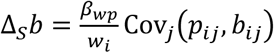and 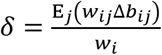, we get the Covariance Based Law of Effect (CLOE):

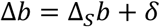

#### 11.2 Estimating Operant Selection from Molar Feedback Functions

For each behavioral measure *b* (which may be either time allocation, peck speed, or peck accuracy), the average change in *C* per unit change in *b* (holding everything else constant) can be expressed by a linear function of the form:

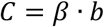

where *β* is the slope of the feedback function. For two different trials 1 and 2, the observed change in behavior (Δ*b*) is defined as:

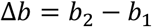

Given the slopes of the feedback functions for the two trials (*β*_1_ and *β*_2_, respectively), the observed change becomes:

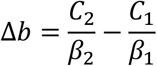

with *C*_1_ and *C*_2_ being the capture rates observed in trials 1 and 2, respectively. In the null-model, the slopes of the two feedback functions remain unchanged, while the capture rate is fixed to the one observed in the first trial. Consequently, the change predicted by the null-model (*δ*) is given by:

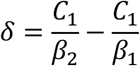

Since the predicted change from the null-model corresponds to the amount of change that would be expected in the absence of selection (i.e., the non-selection term, *δ*, in the CLOE), it follows that the change in behavior due to selection (Δ_S_*b*) can be calculated by taking the difference between the observed and the predicted changes:

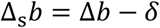

Substituting with the corresponding terms from the above model, we obtain:

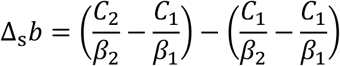

which simplifies to:

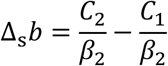

Hence, the amount of behavioral selection may be estimated by taking the difference between the observed value of *b* in the second trial (which is equal to *C*_2_/*β*_2_) and the predicted value of *b* in the second trial (which is equal to *C*_1_/*β*_2_) from the null-model.

## Footnotes

1 Note that statistical fitness predictors are not limited to events that directly increase fitness (such as mating or feeding), but also includes indirect predictors of fitness, such as information about consistent cues for the availability of food, e.g., Anselme (2022); Fortes et al. (2016); McDevitt et al. (2016); Zentall (2016). Parameters that control information seeking might also be relevant predictors of evolutionary fitness, even if their influence may be less direct (e.g., Inglis et al., 1997; cf. also Borgstede, 2021).

2 See Appendix 1 for a mathematical derivation of the CLOE from the elementary Price equation.

3 The analysis would be equally possible for random foraging behavior. In this case, the slope of the feedback function would also depend on the number of food items that have already been retrieved, yielding a negatively accelerated change in expected feedback. Re-analysis of the data under the assumption of random foraging changed the quantitative estimates of the MLBS but did not change the overall qualitative patterns or the group level effects.

4 This identity can easily be verified since, by definition, capture rate may be decomposed such that *R*/*T* = *R*/*P*^+^ · *P*^+^/*T*^+^ · *T*^+^/*T*.

5 A mathematical derivation of the calculations is provided in Appendix 2

6 Since there are nine individuals, there are 2^9^ = 512 possible permutations from which the test distribution is constructed. Consequently, a *p*-value of · 004 means that only two out of 512 permutations deviate at least as much from the null hypothesis (“no change”) as the observed data. For a two-sided test this means that the observed test statistic was the most extreme deviation in the observed direction out of all possible permutations (see Edgington & Onghena, 2007 for a detailed exposition).

## References

Anselme, P. (2022). The optimality of “suboptimal” choice: A psycho-evolutionary perspective. In M. Krause, K. L. Hollis, & M. R. Papini (Eds.), Evolution of Learning and Memory Mechanisms (pp. 193–209). Cambridge University Press.

Anselme, P., Wittek, N., Oeksuez, F., & Güntürkün, O. (2022). Overmatching under food uncertainty in foraging pigeons. Behavioural Processes(201), 104728. 10.1016/j.beproc.2022.104728

Baum, W. M. (1974). On two types of deviation from the matching law: bias and undermatching. Journal of the Experimental Analysis of Behavior, 22(1). 231–242. 10.1901/jeab.1974.22-231

Baum, W. M. (1987). Random and systematic foraging, experimental studies of depletion, and schedules of reinforcement. In A. C. Kamil, J. R. Krebs, & H. R. Pulliam (Eds.), Foraging Behavior (pp. 587–605). Plenum Press.

Baum, W. M. (2012). Rethinking reinforcement: Allocation, induction, and contingency. Journal of the Experimental Analysis of Behavior, 97(1), 101–124. 10.1901/jeab.2012.97-101

Baum, W. M. (2015). The role of induction in operant schedule performance. Behavioural Processes, 114, 26–33. 10.1016/j.beproc.2015.01.006

Baum, W. M. (2017). Selection by consequences, behavioral evolution, and the price equation. Journal of the Experimental Analysis of Behavior, 107(3), 321–342. 10.1002/jeab.256

Baum, W. M. (2018). Three laws of behavior: Allocation, induction, and covariance. Behavior Analysis: Research and Practice, 18(3), 239–251. 10.1037/bar0000104

Baum, W. M. (2023) Introduction to Behavior. An Evolutionary Perspective. Wiley.

Becker, A. M. (2019). The flight of the locus of selection: Some intricate relationships between evolutionary elements. Behavioural Processes, 161, 31–44. 10.1016/j.beproc.2018.01.002

Berlyne, D. E. (1957). Uncertainty and conflict: A point of contact between information-theory and behavior-theory concepts. Psychological Review, 64, Part 1(6), 329–339. 10.1037/h0041135

Borgstede, M. (2020). An evolutionary model of reinforcer value. Behavioural Processes, 104109. 10.1016/j.beproc.2020.104109

Borgstede, M. (2021). Why do individuals seek information? A selectionist perspective. Frontiers in Psychology. 10.3389/fpsyg.2021.684544

Borgstede, M. (2023). Fisher’s fundamental theorem. In T. K. Shackelford (Ed.), Encyclopedia of Sexual Psychology and Behavior. Springer Cham.

Borgstede, M. (2024). Behavioral selection in structured populations. Theory in Biosciences. 10.1007/s12064-024-00413-8

Borgstede, M., & Eggert, F. (2021). The formal foundation of an evolutionary theory of reinforcement. Behavioural Processes, 186, 104370. 10.1016/j.beproc.2021.104370

Borgstede, M., & Eggert, F. (2023b). Squaring the circle: From latent variables to theory-based measurement. Theory & Psychology, 33(1), 118–137. 10.1177/09593543221127985

Borgstede, M., & Luque, V. J. (2021). The covariance based law of effect: A fundamental principle of behavior. Behavior and Philosophy, 49, 63–81. https://behavior.org/wp-content/uploads/2022/01/BP-v49-Borgstede2_corrected.pdf

Broadbent, D. E. (1961). Behaviour. Methuen.

Campbell, D. T. (1956). Adaptive behavior from random response. Behavioral Science, 1(2), 105–110. 10.1002/bs.3830010204

Davison, M., & McCarthy, D. (2016). The Matching Law. Routledge. 10.4324/9781315638911

Donahoe, J. W., Burgos, J. E., & Palmer, D. C. (1993). A selectionist approach to reinforcement. Journal of the Experimental Analysis of Behavior, 60(1), 17–40. 10.1901/jeab.1993.60-17

Donahoe, J. W. (2011). Selectionism. In K. A. Lattal & P. A. Chase (Eds.), Behavior theory and philosophy (Vol. 33, pp. 103–128). New York, London: Springer. 10.1007/978-1-4757-4590-0_6

Edgington, E., & Onghena, P. (2007). Randomization Tests, Fourth Edition (4th ed.). Statistics. CRC Press.

Fortes, I., Vasconcelos, M., & Machado, A. (2016). Testing the boundaries of “paradoxical” predictions: Pigeons do disregard bad news. Journal of Experimental Psychology. Animal Learning and Cognition, 42(4), 336–346. 10.1037/xan0000114

Gilbert, R. M. (1970). Psychology and biology. Canadian Psychologist/Psychologie Canadienne, 11(3), 221–238. 10.1037/h0082574

Hamilton, J. D. (1994). Time series analysis. Princeton Univ. Press.

Hendry, D. P. (1965). Reinforcing Value of Information: NASA Technical Report No. 65-1. University of Maryland.

Herrnstein, R. J. (1961). Relative and absolute strength of response as a function of frequency of reinforcement. Journal of the Experimental Analysis of Behavior, 4, 267–272. 10.1901/jeab.1961.4-267

Herrnstein, R. J. (1964). Will. Proceedings of the American Philosophical Society,, 108(6), 455–458.

Hull, D. L., Langman, R. E., & Glenn, S. S. (2001). A general account of selection: Biology, immunology, and behavior. The Behavioral and Brain Sciences, 24(3), 511–528. 10.1017/S0140525X01004162

Inglis, Forkman, & Lazarus (1997). Free food or earned food? A review and fuzzy model of contrafreeloading. Animal Behaviour, 53(6), 1171–1191. 10.1006/anbe.1996.0320

Kamin, L. J. (1969). Predictability, surprise, attention and conditioning. In B. A. Campbell & R. M. Church (Eds.), Punishment and aversive behavior (pp. 279–296).

Killeen, P. (1972). The matching law. Journal of the Experimental Analysis of Behavior, 17, 489–495. 10.1901/jeab.1972.17-489

Killeen, P. (2023). Theory of reinforcement schedules. Journal of the Experimental Analysis of Behavior. 10.1002/jeab.880

Luque, V. J. (2017). One equation to rule them all: a philosophical analysis of the Price equation. Biology & Philosophy, 32(1), 97–125. 10.1007/s10539-016-9538-y

Luque, V. J., & Baravalle, L. (2021). The mirror of physics: on how the Price equation can unify evolutionary biology. Synthese, 199, 12439–12462. 10.1007/s11229-021-03339-6

McDevitt, M. A., Dunn, R. M., Spetch, M. L., & Ludvig, E. A. (2016). When good news leads to bad choices. Journal of the Experimental Analysis of Behavior, 105(1), 23–40. 10.1002/jeab.192

Okasha, S., & Otsuka, J. (2020). The Price equation and the causal analysis of evolutionary change. Philosophical Transactions of the Royal Society of London. Series B, Biological Sciences, 375(1797), 20190365. 10.1098/rstb.2019.0365

Palmer, D. C., & Donahoe, J. W. (1992). Essentialism and selectionism in cognitive science and behavior analysis. The American Psychologist, 47(11), 1344–1358. 10.1037/0003-066X.47.11.1344

Premack, D., & Premack, A. J. (1963). Increased eating in rats deprived of running. Journal of the Experimental Analysis of Behavior, 6, 209–212. 10.1901/jeab.1963.6-209

Price, G. R. (1970). Selection and Covariance. Nature, 227(5257), 520–521. 10.1038/227520a0

Price, G. R. (1972). Extension of covariance selection mathematics. Annals of Human Genetics, 35(4), 485–490. 10.1111/j.1469-1809.1957.tb01874.x

Pringle, J. (1951). On the Parallel Between Learning and Evolution. Behaviour, 3(1), 174–214. 10.1163/156853951X00269

R Core Team. (2022). R: A Language and Environment for Statistical Computing. https://www.R-project.org/

Simon, C. (2020). The ontogenetic evolution of verbal behavior. European Journal of Behavior Analysis. 10.1080/15021149.2019.1710034

Simon, C., & Hessen, D. O. (2019). Selection as a domain-general evolutionary process. Behavioural Processes, 161, 3–16. 10.1016/j.beproc.2017.12.020

Skinner, B. F. (1966). The phylogeny and ontogeny of behavior. Contingencies of reinforcement throw light on contingencies of survival in the evolution of behavior. Science (New York, N.Y.), 153(3741), 1205–1213. 10.1126/science.153.3741.1205

Skinner, B. F. (1981). Selection by consequences. Science (New York, N.Y.), 213(4507), 501–504.

Staddon, J. E. R. (2020). The Role of Theory in Behavior Analysis: A Response to Unfinished Business,

Travis Thompson’s Review of Staddon’s New Behaviorism (2nd edition). The Psychological Record, 1–7. 10.1007/s40732-020-00409-y

Thorndike, E. L. (1900). The associative processes in animals. Biological Lectures from the Marine Biological Laboratory of Woods Holl, 1899, 69–91.

Timberlake, W., & Allison, J. (1974). Response deprivation: An empirical approach to instrumental performance. Psychological Review, 81(2), 146–164. 10.1037/h0036101

Zentall, T. R. (2016). Resolving the paradox of suboptimal choice. Journal of Experimental Psychology. Animal Learning and Cognition, 42(1), 1–14. 10.1037/xan0000085

